# The instantaneous multi-pronged defense system of latex against general plant enemies

**DOI:** 10.1101/2020.06.19.161869

**Authors:** Luis Francisco Salomé-Abarca, Dejan Gođevac, Min Sun Kim, Geum-Sook Hwang, Sang Cheol Park, Young Pyo Jang, A. M. J. J. van den Hondel Cees, Robert Verpoorte, Peter G. L. Klinkhamer, Young Hae Choi

## Abstract

Based on the hypothesis that variation of the metabolomes of latex is a response to selective pressure and should thus be affected differently from other organs, their variation could provide insight into the defensive chemical selection of plants. Metabolic profiling was utilized to compare tissues of *Euphorbia* species collected in various regions. The metabolic variation of latexes was much more limited than that of other organs. In all of the species, the levels of polyisoprenes and terpenoids were found to be much higher in latexes than in leaves and roots. Polyisoprenes were also observed to physically delay the contact and growth of pathogens with plant tissues. A secondary barrier composed of terpenes and, in particular, 24-methylenecycloartanol, exhibited antifungal activity. These results, together with the known roles of the enzymes also present in latexes, demonstrate that they are part of a cooperative defense system that comprises both biochemical and physical elements.

## Introduction

Living organisms are constantly challenged by diverse environmental factors, including biotic and abiotic^1^. In order to survive as a species, they must develop responses during these interactions, evolving genetically and/or phenotypically (Andrew et al., 2010). Therefore, the biological and chemical complexity of life can be considered to be, to some degree, the consequence of genetic diversity modulated by ecological interactions. Especially in plants, sessility has acted as a strong evolutionary pressure, resulting in the development of diverse physical and biochemical tools as sophisticated adaptive systems. This has allowed them to endure throughout time in diverse ecosystems (Bucharova et al., 2016). Among these tools, plant exudates have attracted great interest due to their adaptive origin, having resulted from their co-evolution with other organisms, including insect herbivores and microorganisms (Konno 2011). Plant exudates represent one of the surface defense layers associated with both primary and secondary defenses. Among plant exudates, latexes have attracted much focus, not only because of their commercial value, but also due to their distinctive chemistry, which is assumed to possess ecological potential as a primary barrier in response to exogenous factors (Agrawal and Konno, 2009).

Although the specific role of individual latex metabolites remains unknown, as a whole, their metabolomes are highly distinctive, exhibiting chemical fingerprints that differ both quantitatively and qualitatively from those of other plant organs (Seiber et al., 1982; Konno et al., 2004; Konno et al., 2006). Furthermore, the bioactive metabolite contents of latex at damaged points before and upon an attack can vary, as their concentration is locally increased (Ball et al., 1997; Hölscher et al., 2016; Gorpenchenko et al., 2019). Therefore, elucidating the roles of latex metabolites and their functions in the complex mechanisms of plant defense systems could contribute to a greater understanding of chemical selection in the evolution of plant defense. Together with chemical selection, the degree of metabolic variation in defensive exudates, such as latexes, could also determine the success of this type of defensive system. As an approach to understand the detailed roles and mechanisms behind latex chemistry and its variations, a method was designed to answer the following fundamental questions: are latexes from different species chemically similar? To what extent do environmental factors affect the chemical variation of latexes? Do specialized metabolites exert their biological activity individually or do they act in a complementary, or even synergistic, manner with other latex components? If so, how do they interact?

Considering the complexity of factors involved in the issues to be investigated, a holistic approach was adopted. The experiments were performed using a model system consisting of a plant sample set with different genetic (species) and environmental (geographical origin) backgrounds. The sample set contained wild *Euphorbia palustris*, *Euphorbia amigdaloides*, and *Euphorbia glareosa* (Euphorbiaceae) specimens collected in different locations in Serbia. The metabolic composition of latexes, leaves, and roots were studied by ^1^H NMR and LC-MS-based metabolomics, and GC-MS and HPTLC-DART-MS were utilized as supplementary tools for analysis of targeted metabolites. Based on the metabolomics results, the anti-herbivory, antibacterial, and antifungal activities of the latexes were assayed, and the degree of chemical and biological variation was correlated in order to determine the role of individual metabolites. This holistic approach revealed the potential of latexes to play an efficient role in the defense system of plants.

## Results

### Metabolic variation in *Euphorbia* latexes by geographical locations; latexes have less metabolic variation than other tissues

To investigate and compare the effect of environmental factors on the metabolome of latex with that observed in other organs (leaves and roots), three *Euphorbia* species were collected from nine locations in Serbia. The underlying concept was to gauge similarity in the metabolic variations of latexes and tissues from the plant samples. A lower variation in latexes, as compared to the tissues, could imply that they are involved in basic functions, for example, as a first barrier against general predators or pathogens. Conversely, if the variation was greater, it could indicate that latexes are related to specialized roles, such as a reaction to subtle environmental changes, including attacks of location-specific predators or pathogens.

To compare the metabolic variation of each organ and latex, their ^1^H NMR spectral data were subjected to multivariate data analysis (MVDA). Firstly, the spectra of the samples from the three locations were analyzed by principal component analysis (PCA). As shown in Supplementary Figure 1, the metabolome of all of the profiled samples was distinctive for each *Euphorbia* species. This was not unexpected, as the species is known to have a significant effect on the metabolome. Interestingly, however, when comparing the metabolic variation related to the geographical origin of leaves, roots and latexes, the latter exhibited much less variation than the others. This was also confirmed by further MVDA, a soft independent modelling of class analogy (SIMCA) analysis, in which a PCA model is built for each class, and the distance (DModX) between models is measured to determine their similarity or dissimilarity (Eriksson et al., 2011).

For all of the species, the values of DModX based on their geographical origin were calculated for latexes, leaves and roots, and the obtained values were logarithmic-transformed and averaged to identify the degree of variation. In this model, the higher was the DModX, the higher was the metabolic variation. As shown in Figure 1, the SIMCA model verified that the variation in metabolomes among latexes was much lower than that between leaves and roots for all of the tested species.

**Figure 1.**
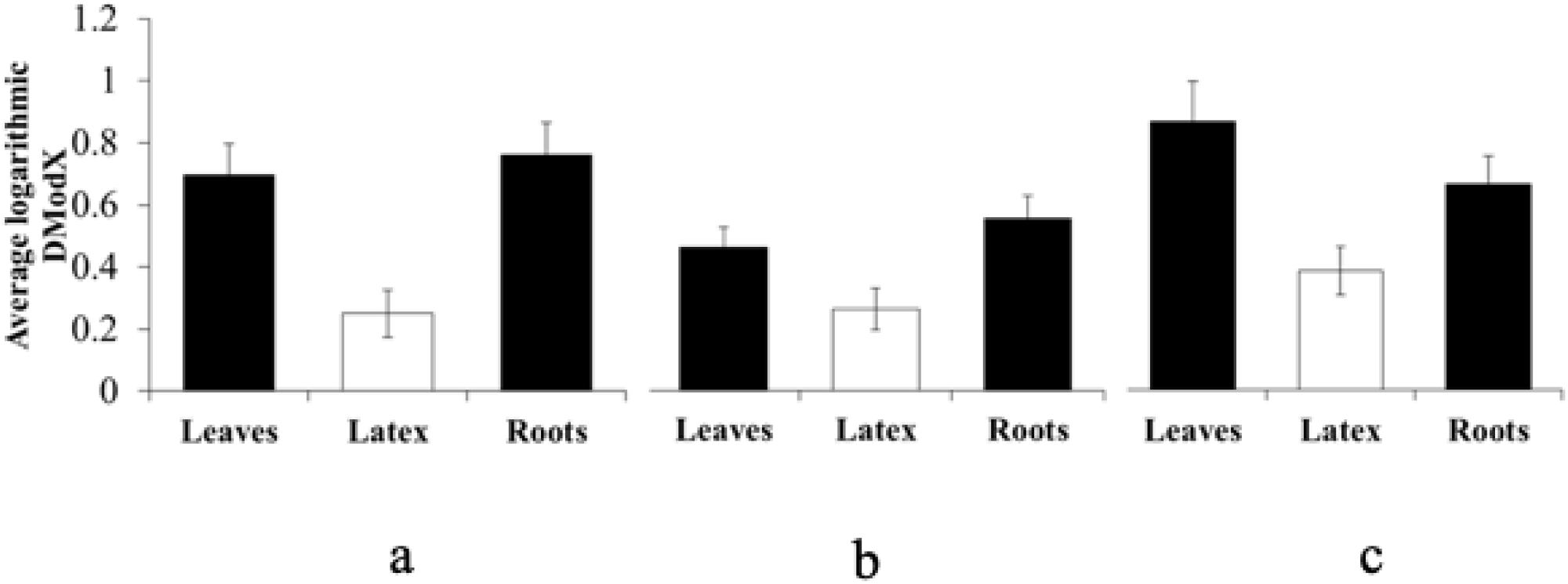
Logarithmic averaged distance to the model (DModX) values in latex, leaves, and roots of three Euphorbia species. A, *Euphorbia amygdaloides.* B, *Euphorbia glareosa*. C, *Euphorbia palustris*. The averaged values represent the mean (n = 30 ± standard error) of the DModX from a soft independent model of class analogy (SIMCA) analysis of all samples of leaves, roots, and latex per species.

To confirm this, the *Q^2^* value (an indicator of correlation) of partial least square-discriminant analysis (PLS-DA) was calculated. The *Q^2^* value reflects the degree of correlation between the chemical data set and the classes (Eriksson et al., 2011) (geographical origins), and can thus be regarded as a measurement of their degree of total correlation. Consequently, a lower degree of correlation would be anticipated between the latex chemical set and its geographical origins. As shown both by PCA and SIMCA results, geographical factors exert highly differential effects on the metabolic variation of leaves (*Q^2^* = 0.96 ± 0.002) and roots (*Q^2^* = 0.91 ± 0.012), while latexes showed much less correlation (*Q^2^* = 0.53 ± 0.220), indicating that they are not affected as much by their geographical origin as the tissues.

All of the data analyses confirmed that the metabolome of latexes is more constant than those of leaves and roots. They are thus apparently less influenced by environmental factors, being possibly involved in generic defense responses, for example, acting as a first barrier against general enemies. If this is the case, it could also be supposed that latexes could be less influenced by the species factor than by geographical locations. Among the tested species, the metabolic variation in latexes (DModX = 0.76 ± 0.07) associated with species factors was lower than in roots (DModX = 1.05 ± 0.08), but showed no relevant difference with that of leaves (DModX = 0.76 ± 0.06).

The next step consisted of identification of the discriminating metabolites between latexes and other tissues. This was carried out using their ^1^H NMR spectra. The lower chemical variation observed in latexes indicates that there are more common metabolites among latexes than in leaves and roots. The comparison of the ^1^H NMR spectra of leaves, roots, and latexes revealed extremely higher levels of triterpenes in latexes than in the other tissues. Polyisoprenes, in particular, were present in high amounts in latexes, as compared to leaves and roots in all of the studied *Euphorbia* species. The latex-specific triterpenes were characterized by methyl signals in the δ 0.70 – δ 1.90 range (Figure 2). Additionally, the cyclopropane moiety present in the structure of cycloartanol was identified by two doublets at δ 0.55 (d, *J* = 4.0 Hz) – δ 0.35 (d, *J* = 4.0 Hz) assigned to the 19-*endo* and 19-*exo* protons, respectively (Figure 3). The cycloartanols were further confirmed by DART-MS and GC-MS analysis. The direct mass analysis showed 441.4167 *m/z* and 458.4429 *m/z*, which were assigned to the [M+H]^+^ and [M+NH_4_]^+^ adducts (mass error < 7 ppm), respectively, of 24-methylenecycloartanol. In terms of the relative quantitation performed using integration of the H-19 resonance, it was found that the content of the cycloartanol-type triterpenes in latex was over eight times higher than in other tissues (Figure 3), comprising almost 16% of the dry weight of latexes. The GC-MS analysis confirmed this, revealing that a few other cycloartane analogues, such as 24-methylenecycloartanone and 24-methylenecycloartanol acetate, were more abundant in latexes than in other organs.

**Figure 2.**
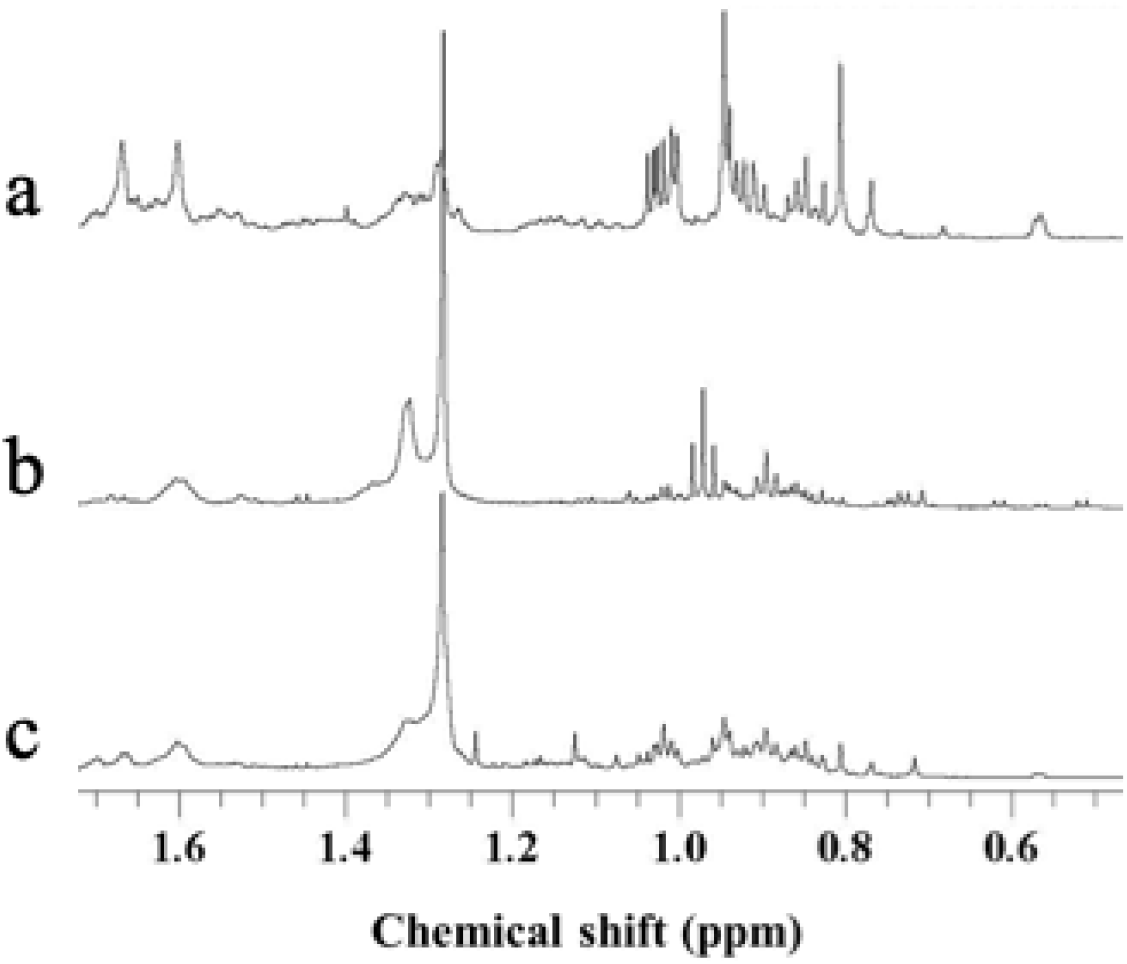
Representative ^1^H NMR spectra (600 MHz, CH_3_OH-*d_4_*) from latex, leaves, and roots. A, latex. B, leaves. C, roots. Range of δ 0.4 – δ 1.8 of latex reveals a higher concentration of terpenes resonances.

**Figure 3.**
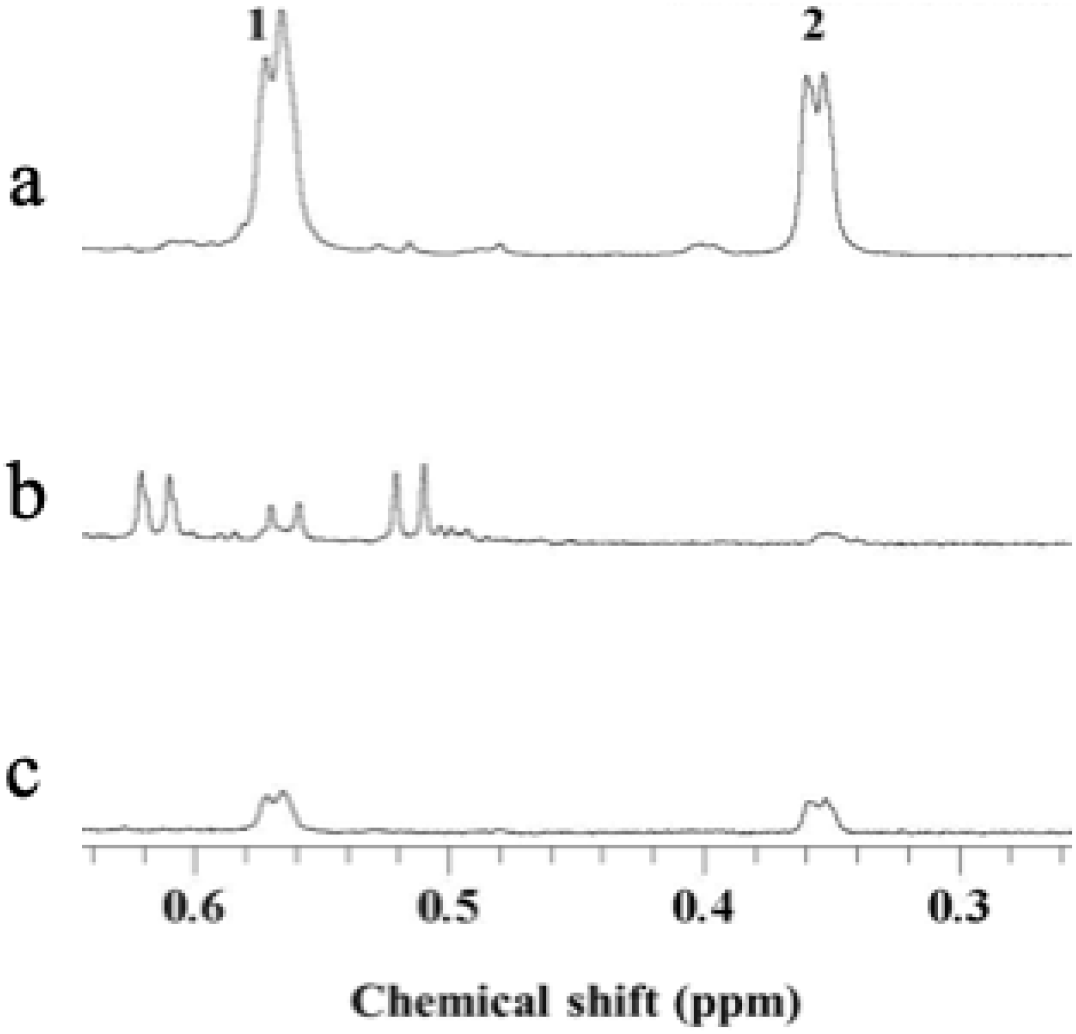
Expanded ^1^H NMR spectra (600 MHz, CH_3_OH-*d_4_*) from latex, leaves, and roots. A, latex. B, leaves, C, roots for cyclopropane group. Two doublets at (1) δ 0.55 and (2) δ 0.35 correspond to the 19-*endo* and 19-*exo* protons, respectively, of the cyclopropane ring of 24-methylenecycloartanol with a concentration approximately eight times higher in latexes than in the other two tissues.

The LC-MS analysis of the latex samples provided further details of their metabolic variation. Similarly to the ^1^H NMR-based analysis, variation among latex samples from different locations was found to be low, and *E. palustris* and *E. glareosa* showed no separation by geographical origin. However, some species-specific metabolites, such as alkaloids, were detected (Table 1). These alkaloids were found to be specific for *E. glareosa* and *E.amygdaloides*, while acyl sugars were determined to be more related to *E. palustris*.

**Table 1.**
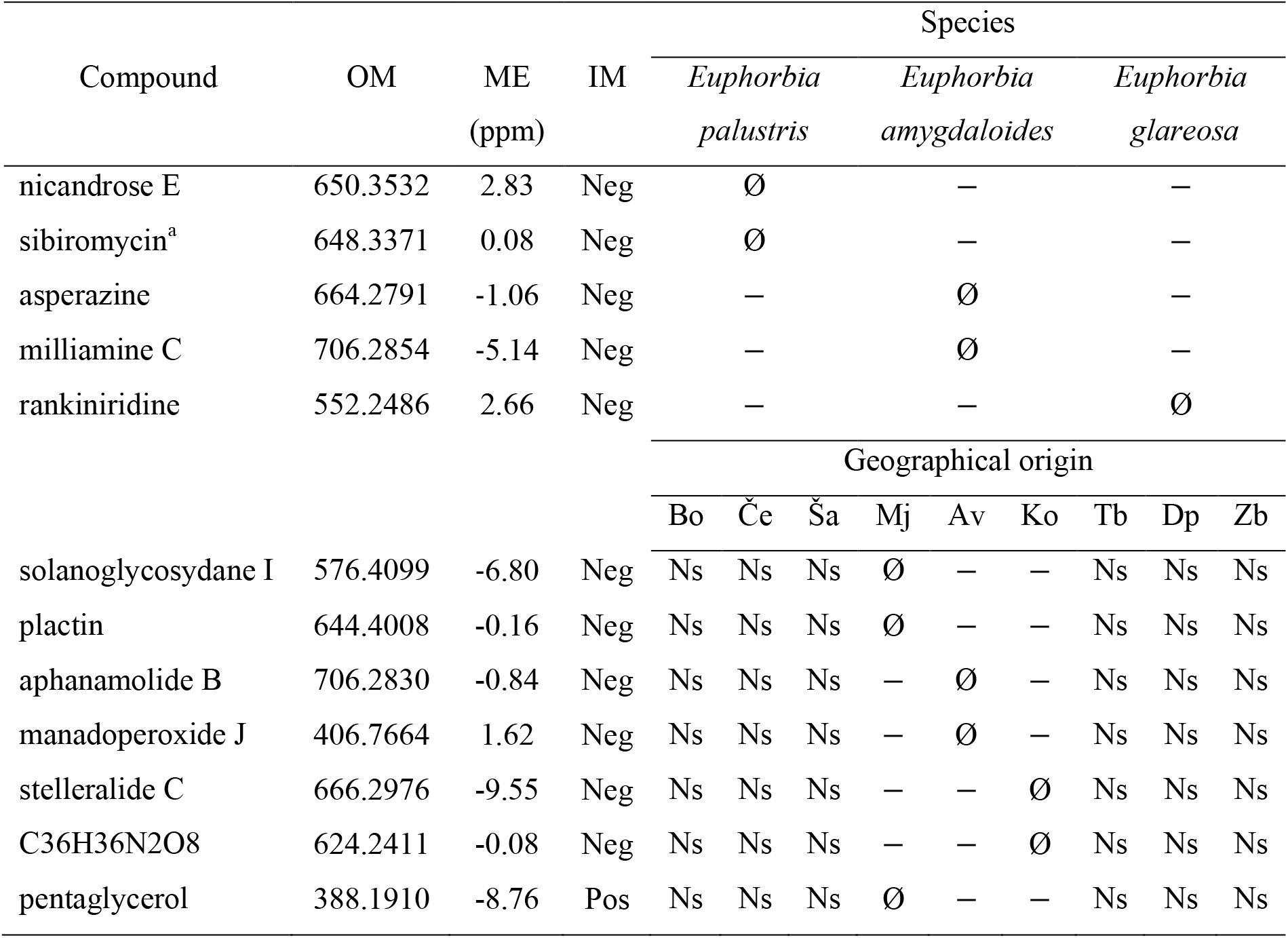
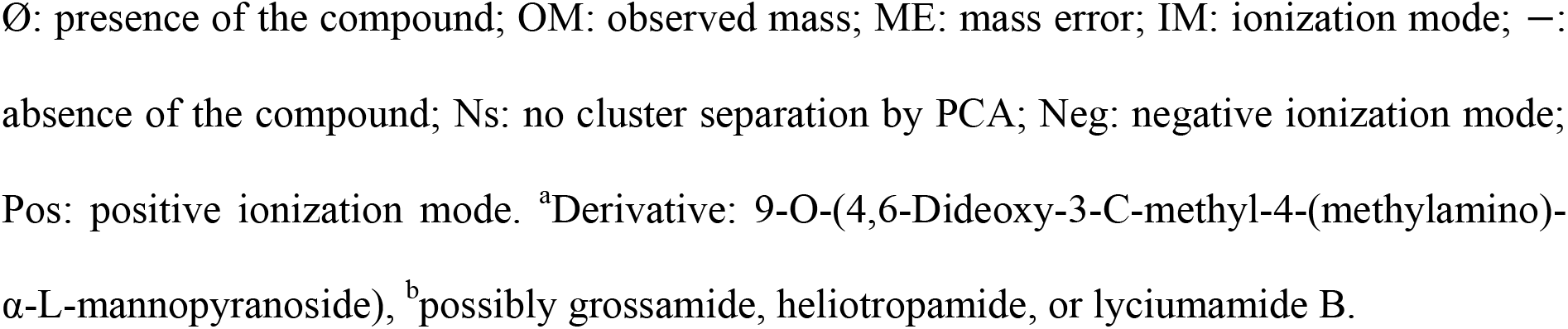
Identified discriminant metabolites in *Euphorbia palustris, Euphorbia amygdaloides*, and *Euphorbia glareosa* latex collected in different locations of Serbia.

### Latex metabolites have more stable anti-herbivory activity

Latex is believed to play a primary role in the defense of plants against a broad range of natural enemies, acting instantly upon attack as a first barrier while other induced defense mechanisms are being activated. As a consequence, if the variation in the chemical composition of latexes is lower than in other tissues, the variation in their anti-herbivore activity should also be lower than that of the other tissues. To prove this hypothesis, latexes, leaf, and root extracts were challenged with a generalist herbivore, *Mamestra brassicae*. In terms of anti-herbivory activity, the results showed significant anti-feeding effects for all three types of extracts for two of the three tested species compared to the negative control diet (*p* < 0.05). In the case of *E. palustris*, the latex exhibited a lower, but statically insignificant (*p =* 0.12), average medium than the negative control. This value, however, denotes a trend towards positive anti-herbivory activity. In addition, when analyzing the variation provided by the standard error of the bioactivity for each species, the variation of the anti-herbivore activity in its latex was lower than that of leaves and roots at least in two of the three studied species (Figure 4). This result suggested that the lower chemical variation of latexes was consistent with more homogeneous variation in biological activity against *M. brassicae*.

**Figure 4.**
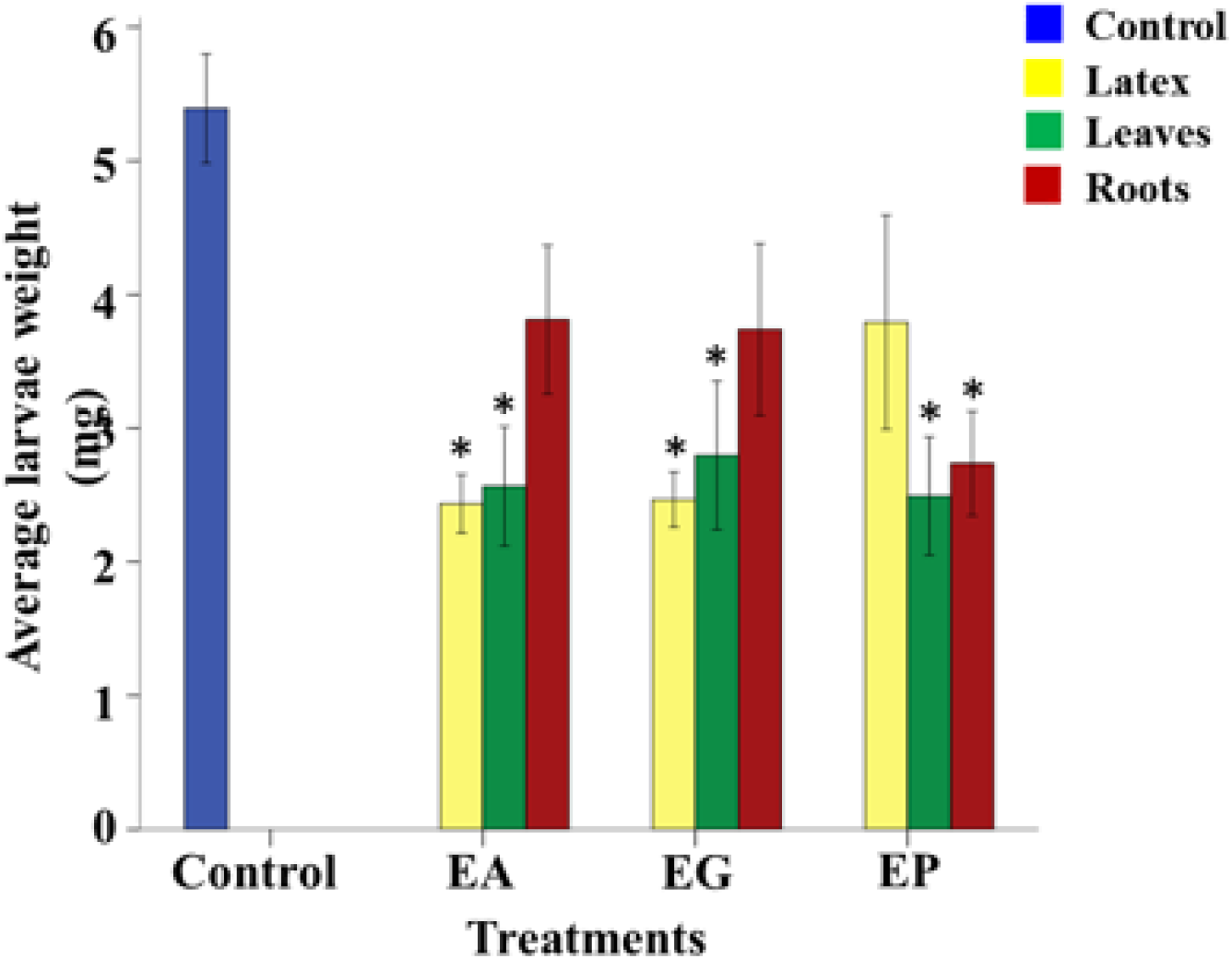
Anti-herbivore activity from latex, leaves, and roots of *Euphorbia amygdaloides*, *Euphorbia glareosa*, and *Euphorbia palustris*. The values represent the mean value (n = 30) ± standard error. * represents significant differences between the treatment and the control in a least-square means test (α = 0.05). EA, *Euphorbia amygdaloides*, EG, *Euphorbia glareosa*, EP, *Euphorbia palustris*.

### Latex, as a plant defense against microorganisms, is both chemical and physical

If latexes are involved in a basic defense mechanism, they should also provide protection against pathogenic bacteria or fungi. However, when latex extracts were tested on *Pseudomonas viridiflava*, *Pseudomonas fluorescens* and *Pseudomonas putida*, considered to be general pathogens, they did not exhibit any antibacterial activity. A possible explanation for these results was that latexes may provide a physical or mechanical defense mechanism rather than, or added to, some chemical toxicity, given the high concentrations of polyisoprenes detected with metabolic screening. This assumption fits well with the hypothesis that latexes have a general and broad function of defense in the early stage of plant defense. To test this hypothesis, a mimicking experiment was carried out. A layer of poly-*cis*-1,4-isoprene was spread on agar plates inoculated with the bacteria, and their movement was observed, i.e., to determine whether or not they could move through the rubber layer. All of the tested bacteria were unable to move through the rubber layer (Figure 5) demonstrating, for the first time, that the mechanical defense of latexes could *per se* suffice to defend the plants from bacterial infections, or at least, that the barrier could delay their movement, and thus the onset of an infection.

**Figure 5.**
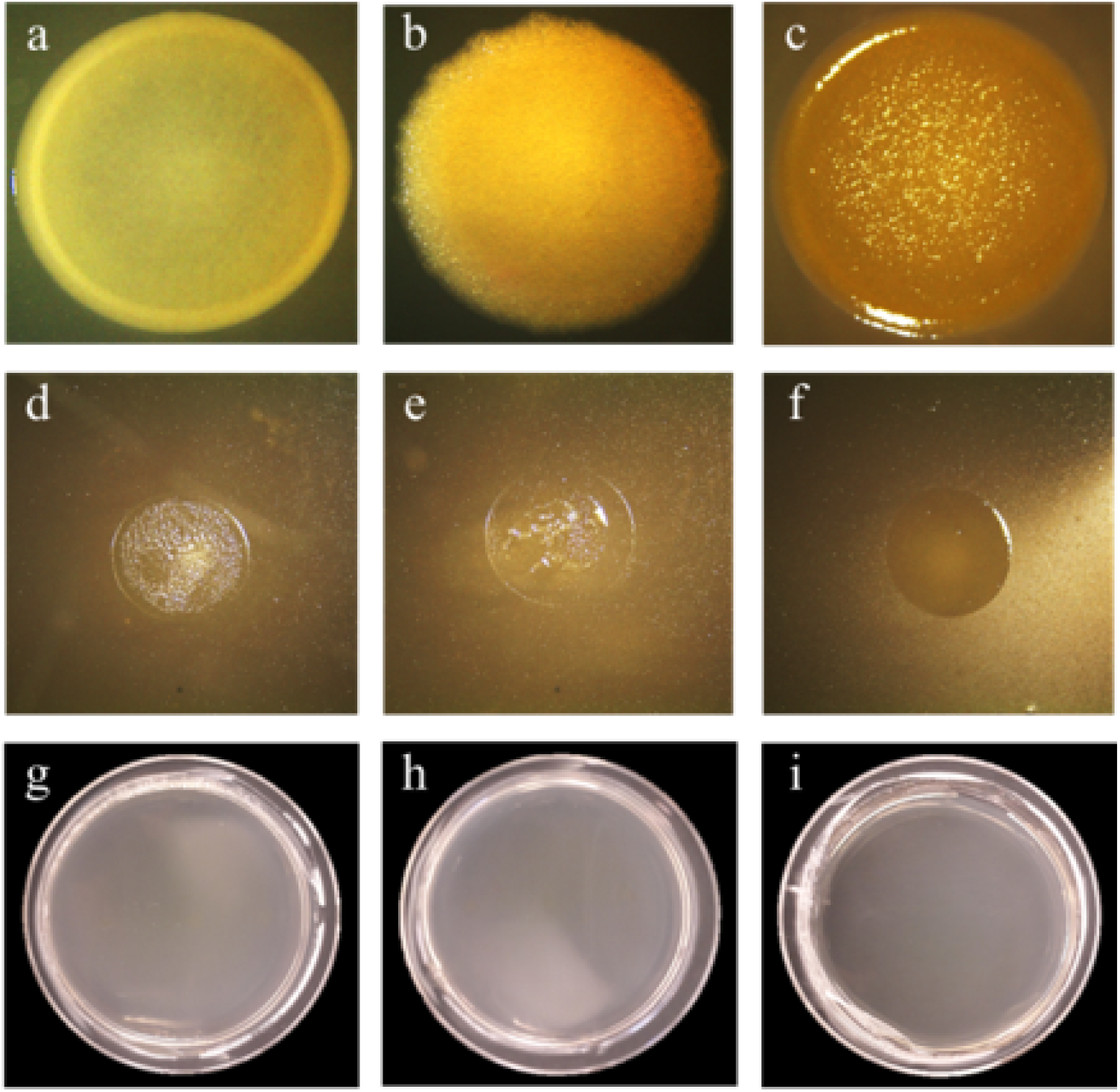
Antibacterial effect of rubber against bacterial pathogens. A, *Pseudomonas fluorescens*. B, *Pseudomonas putida*. C, *Pseudomonas viridiflava*. The first three images represent the negative control consisting of the bacteria growth in Mueller-Hinton agar (nutrient agar 2 for *P. viridiflava*) after 24 h; D, E, and F: the same bacteria growing on top of the rubber layer. The bacteria were not able to penetrate the layer and grow in the agar media; G, H, and I: growth control for the rubber treatment. The rubber layer with bacteria above it was removed, and the plates were incubated for 24 h to reveal possible bacterial growth.

In the case of fungi (*Botritys cinerea* and *Alternaria alternata*), a similar phenomenon, although with slight differences, was observed. When testing the previously described sealing effect, the layer did not indefinitely block the spread of fungal growth and, after some time, the fungi were able to penetrate it. However, movement was definitely delayed, so that when the fungi were placed on top of the rubber layer, they were eventually able to break through it, but at a very slow rate. Moreover, when the diameter of the colony submitted to the rubber treatment was compared to the positive control, radial growth was significantly reduced (Figure 6a). A possible complementation between polyisoprenes and small molecules in latex was assessed by supplementing the rubber slurry with latex. Results revealed a significant decrease in radial fungal growth compared to that observed with rubber alone (Figure 6b).

**Figure 6.**
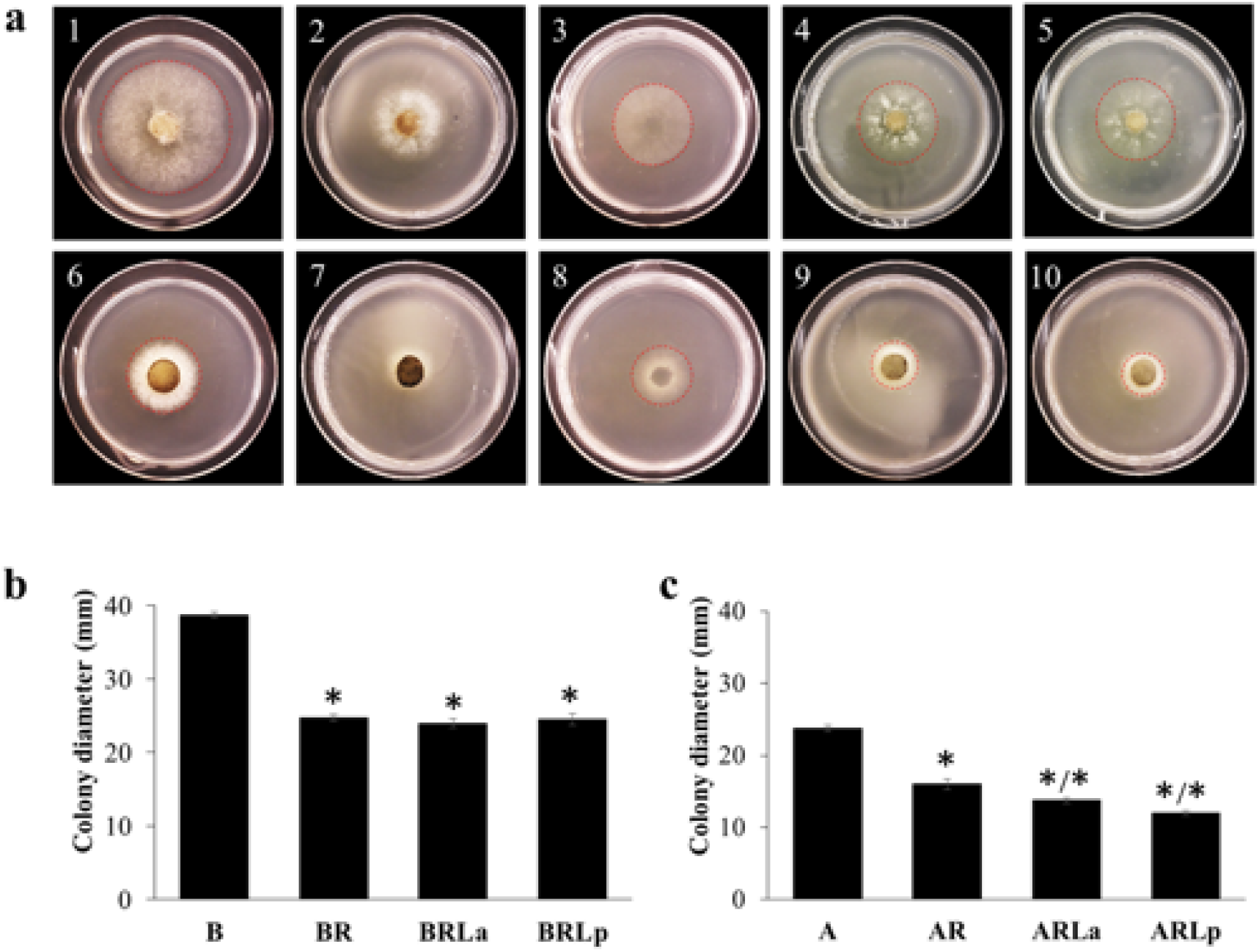
Antifungal activity of rubber and rubber plus latex against *Botrytis cinerea* and *Alternaria alternata*. A, Radial growth of *Botrytis cinerea* and *Alternaria alternata* in control and treated medium at 48 h. 1, Growth control of *B. cinerea* growing on PDA medium. 2, *B. cinerea* growing on PDA medium covered with a rubber layer. 3, Mycelium of *B. cinerea* in the agar medium after removing the rubber layer. 4, *B. cinerea* growing on PDA medium covered with a rubber layer combined with 200 μg/mL of *E. palustris* latex. 5, *B. cinerea* growing on PDA medium covered with a rubber layer combined with 200 μg/mL of *E. glareosa* latex. 6 – 10 represent the same treatment order, but against *Alternaria alternate.* B, Antifungal activity of rubber and rubber supplemented with latex 48 h against *B. cinerea*. The values represent the mean (n = 4) ± standard error. The stars represent significant differences between the treatment and the control (p ≤ 0.05) in a Dunnett test. B: growth control of *Botrytis cinerea*. BR: *B. cinerea* growing medium covered with a rubber layer. BRLa, *B. cinerea* growing medium covered with a rubber layer supplemented with latex of *E. amygdaloides*. BRLp, *B. cinerea* growing medium covered with a rubber layer supplemented with latex of *E. palustris*. C, Antifungal activity of rubber and rubber supplemented with latex at 48 h against *A. alternata*. The order of treatments is the same as for *B. cinera*, but A means *Alternaria alternata*. For example, AR: *A. alternata* growing medium covered with a rubber layer. The values represent the mean (n = 4) ± standard error. The stars represent significant differences between the treatment and the control, and a star after a slash indicates significant differences between the treatment and B/AR (*p* ≤ 0.05) in a Dunnett test. Fungal pathogens were able to penetrate the mechanical barrier of rubber, but their proliferation was significantly delayed.

### The role of triterpenoids selectively accumulated in latexes: fungicidal effect against residual fungi beyond the polyisoprene barrier

In the metabolomics analysis, all of the tested *Euphorbia* latexes collected from various locations contained a very high amount of cycloartanols together with polyisoprenes. Unlike bacteria, the tested fungi were eventually able to penetrate the rubber barrier, albeit after an initial delay (Figure 6a). Being the major and selectively concentrated metabolites in latex, a cycloartanol isolated from *Euphorbia* latexes, 24-methylenecycloartanol, was chosen as an example and tested for its fungicidal activity and compared with that of other common triterpenes and steroids that are commonly found in leaves and roots.

The minimum inhibitory concentration (MIC) values of 24-methylenecycloartanol and other common steroids and terpenoids, including *β*-sitosterol, *α*-amyrin, *β*-amyrin, ursolic and oleanolic, were determined against *B. cinerea*. With the exception of latex-specific 24-methylenecycloartanol, none of the compounds showed inhibition at the tested concentrations. On the contrary, some of the common steroids or triterpenoids displayed a growth-promoting effect on the fungus. For example, ursolic acid exhibited a strong growth-promotion effect in liquid suspensions of fungi 64 h after inoculation (Supplementary Figure 2). This indicated that 24-methylenecycloartanol plays a specific role in latex. Interestingly, although the isolated compound displayed mild inhibition against a range of concentrations of *B. cinerea* of 500–1000 μg/mL, this occurred in concentrations as high as 11000 μg/mL in latexes.

### A possible combination effect between polyisoprene and cycloartanols: polyisoprene acts as a dispersive agent for cycloartanols efficient activity

The MIC value of 24-methylenecycloartanol is too high for it to be considered truly active as a fungicide. This low activity could be explained partially by its low solubility in the aqueous conditions of the test. As a result, it seemed likely that some component/s in the chemical matrix of the latexes acted as a solubilizing agent. To test this, two aqueous mixtures, one containing 20 mg of 24-methylenecycloartanol alone and another with added poly-*cis*-1,4-isoprene, were prepared and then dried. When the sample containing only the cycloartanol was dried, the compound crystalized; whereas, the mixture with polyisoprene formed a layer coating the wall of the tube, and the sterol was determined to be evenly distributed throughout the layer. This experimental result suggested the potential of rubber as a dispersing agent for lipophilic metabolites, in this case allowing distribution of cycloartanol in latexes to increase its efficiency as a defense mechanism.

## Discussion

Plants are thought to have developed their own defense systems differently to other living beings. However, the detailed mechanisms behind these systems are largely unknown. Plant latexes have long been considered to be highly valuable substances. This is mainly because they are a rich source of selected chemicals, such as natural rubber (1,4-*cis*-polyisoprene) with its important industrial applications, but also due to their potential use as a high hydrocarbon sink for fuel and as chemical feedstocks. Further studies, however, have revealed the presence of bioactive metabolites and proteins with insecticidal, deterrent, antibacterial, antifungal and cytotoxic activity, pointing to a probable role in the mechanisms behind plant defense. Moreover, latexes have also recently been recognized as a valid ecological model for the understanding of plant-herbivore interactions (Konno 2011).

The chemical composition of latex includes three major categories of compounds: polyisoprene polymers (rubber), proteins, and small specialized metabolites. Among these, proteins have been relatively more studied due to their evident biological functions against pathogens and herbivores. In the case of rubber, research has been focused on the coagulation process necessary for industrial purposes, but little is known about its role in defense mechanisms. The chemical and biological diversity of latexes lies in the content of specialized metabolites, similarly to other tissues. However, despite their proven biological activities, the potential roles of these specialized metabolites *per se* or eventually in cooperative roles with other components of latexes, such as rubbers or proteins, related to plant response to the environment have not yet been investigated.

The experimental results obtained provided answers to the fundamental research questions considered to be essential to elucidate the roles and mechanisms of latex in the defense system. First, the need to determine whether the composition of latex is susceptible to environmental conditions was crucial, since it could provide insight into the type of defensive action that could be provided by latex. This is because, due to their anatomical location, they are very likely one of the first mechanical and chemical barriers of plants against herbivores and microorganisms. In this case, latexes should possess distinctive metabolomes from other plant organs. Furthermore, to be able to fulfil their basic role as a first barrier, the latex metabolome should be composed of a few selected metabolites at high concentrations, exhibiting a semi-conserved constitutive chemical composition, and relatively unaffected by environmental factors. To evaluate this possibility, a systemic approach examining, for example, latexes from plants from diverse locations, was thought to useful to provide insight into the basic roles and mechanisms of latex compounds.

Accordingly, the latexes of three latex-bearing species of *Euphorbia*, collected in nine geographical locations, were metabolically characterized. The degree of metabolic variations of latexes, leaves, and roots of the plants was analyzed using several MVDA methods, including PCA, SIMCA, and PLS-DA. From the analyses, it was found that the metabolites in latexes were clearly distinguished from other organs, both qualitatively and quantitatively. In addition, latexes exhibited considerably less metabolic diversity, resulting in lower variation per location than in other organs. It is generally believed that plant metabolomes are largely influenced by environmental factors, and the difference in chemical diversity in ecotypes has been well documented for a few species, and interpreted as an adaptive response to specific biotic and abiotic factors found in different ecosystems (Shelton 2004). Moreover, each plant species evolves differentiated chemical responses according to its survival requirements both at a population and individual levels. Contrary to this general rule, however, in the case of latexes, their metabolome was shown to vary much less than other organs. From a plant defense perspective, this lower metabolic variation of latexes indicates that they might play a primary and general role in defense, anticipating the more complex inducible response.

Among the selectively secreted chemical classes in latexes, triterpenes, specifically cycloartanols, such as 24-methylenecyloartanol, proved to accumulate in latex rather than in leaves and roots. The same applied for polyisoprenes. Triterpenoids have been identified in *Euphorbia* species due to their chemotaxonomic importance (Ponsinet and Ourisson, 1968; Mahlberg and Pleszczynska, 1983), and many of them have been suggested to be chemomarkers for certain taxons (Mahlberg et al., 1987; Mahlberg et al., 1988).

Previous reports on latex chemical composition identify rubber and triterpenes as the main components of rubbery latexes^3,4^ assuming that, given their proximity in biosynthetic pathways, their coexistence in laticifers might have occurred during evolution (Nemethy et al., 1983; Mahlberg 1985; Piazza et al., 1987a; Piazza et al., 1987b;). This hypothesis, however, does not explain the selection of a specific terpenoid in a concentration approximately eight times higher than in other organs, as observed in our results. Indeed, the value of triterpenes in latexes should be functionally coupled to that of rubber. In other words, the lower degree of chemical variation in latexes strongly suggests that their biological activities are limited only to basic protection in an early stage of defense, rather than to a specialized role.

To prove this hypothesis, the anti-herbivory activities of latexes and other organs were measured. The three types of samples (latexes, leaves, and roots) exhibited almost the same degree of anti-herbivory effects against *M. brassicae*, despite possessing largely differentiated metabolomes. In the case of the latex of *E. palustris*, the data of two samples, one from Borča and one from Čenta, constituted outliers for that group of samples, resulting in a large standard error, and probable responsibility for the lack of a significant difference with the negative control. Nonetheless, even with a more limited number of metabolites than the other investigated organs, latexes exhibited a very efficient metabolome designed as a defense against herbivores and pathogens. Additionally, the biological activity of latexes against *M. brassicae* showed less variation among the latex samples, irrespective of their geographical location. Consequently, instead of a disadvantage, the low variation and relatively simple metabolome proved to be able to provide more stable and consistent protection against herbivores than those of leaves and roots, which are more susceptible to environmental variations, as they adapt to them through adjustments in their metabolome. These results point to a long co-evolution between the plant species and diverse herbivores to develop a selective or limited number of compounds against a broad range of natural enemies (Firn and Jones, 2003). In particular, this conserved metabolome could be more beneficial for a primary role in defense when dealing with generalist organisms, combining constitutive defense compounds and mechanical traits. In this regard, it is well-known that when high herbivore pressure conditions are present, in the case of latex-bearing species, it was found that the constitutive defenses could create a cost-benefit balance in plant fitness (Andrew et al., 2007; Moore et al., 2014).

In nature, the most efficient way to deal with a wide range of natural enemies and to avoid the fast development of resistance with a minimal use of resources is to employ a synergistic blend of traits (Richards et al., 2010; Richards et al., 2012). If this is the case for latexes, a possible synergistic effect between their ingredients is to be expected. To examine this potential cooperative mechanism, the activity of polyisoprene was firstly tested against bacteria and fungi. Rubber was found to block the invasion of bacteria completely. In the case of fungi, however, it worked only partially, producing retardation in the growth of the pathogenic fungi. It was thus assumed that metabolites assisted the rubber barrier. Presumably, the hyphae of fungi that survived after penetrating the mechanical rubber barrier would then come in direct contact with latex inside of the laticifer, which contains higher concentrations of active metabolites than the coagulated layer. This is where 24-methylenecycloartanol, by far the most abundant compound in the latexes, could act on the fungi. The fungicidal activity of the cycloartanol was tested against *B. cinerea*, and compared with other common terpenoids and steroids that exist in leaves or roots. Of the tested steroid and terpenoids, only 24-methylenecycloartanol showed fungicidal activity against *B. cinerea*. These results reveal the existence of a systemic defense mechanism with a synergistic effect between rubber and cycloartanol, and highlight how this enables latexes to deal with the threat of a large number of enemies with a very limited number of compounds. Rubber is able to protect plants from a broad range of pathogens and herbivories simply due to a mechanical effect which is complemented with a chemical defense in the form of specific compounds, such as cycloartanols, which act on residual herbivores or pathogens. Moreover, below the film of coagulated rubber, a high concentration of bioactive metabolites produced by induction will accumulate as a response to aggression (Krstić et al., 2016), thus combining mechanical and chemical effects in one single barrier.

Another aspect of the synergistic mechanism involving cycloartanol and rubber is that rubber could function as a natural dispersive agent for the triterpenoids in latex. The results from the tests performed to demonstrate this showed that the triterpene-rubber dispersion and rubber films and their combination with latexes acted as a well-designed system against bacterial and fungal microorganisms. In this system, the rubber particles functioned as a carrier and disperser of phytosterols and triterpenes during latex exudation and throughout the whole production of a sealing film after coagulation. Therefore, the complementation between the mechanical defense (rubber coagulation) and chemical defense (specialized metabolites) resulted in a polymeric film, in which embedded metabolites are evenly distributed in the whole film to increase its effectiveness over the protected area. In addition to the fungicidal activity and function as a dispersion agent, triterpenes might have other functions. They could contribute to the polymeric structure of polyisoprene to strengthen the defensive barrier, as may be deduced from other related experiments (Mironenko et al., 2016). Further work is needed to understand the specificity of the synergy between latex and cycloartanol and, in particular, among many other available steroids and triterpenoids.

This finding denotes the complementation of mechanical and chemical defense systems against herbivories, bacteria, and fungi. The results also indicate that the basic defense system of latexes is attributable to mechanical features combined with selectively secreted metabolites, all of which hardly vary when exposed to environmental changes. Even if this were insufficient to avert fungal infections or fully deter herbivory, the primary latex barrier provides immediate protection until specific (inducible) defense mechanisms that are being activated can exert their specific effect. As an advantage over normal induced responses, the defense system in latexes is much faster, producing a build-up of active compounds in just a couple of minutes (Agrawal and Konno, 2009; Konno 2011).

Based on the experiments in this study, plant latexes are proposed to play a role as a general primary defense using mechanical (e.g., stickiness and coagulation), chemical (constitutive metabolites), and biochemical responses such as specific enzymes (Agrawal and Konno, 2009; Konno 2011; Salomé-Abarca et al., 2019) in a coordinated manner (Figure 7). This kind of systemic primary defense barrier of latexes was proven by characterizing the full latex metabolome, which appears to be quite stable and independent of environmental changes. This was also verified by the higher similarity of the chemical profile of latexes among different species than that of leaves and roots. This similarity was reflected as a lower variation in their composition throughout geographical regions, and indicates semi-conserved constitutive chemical selection in these *Euphorbia* species. This consistency of latex components suggests that it plays a complementary role in the plant defense system in conjunction with other mechanisms.

**Figure 7.**
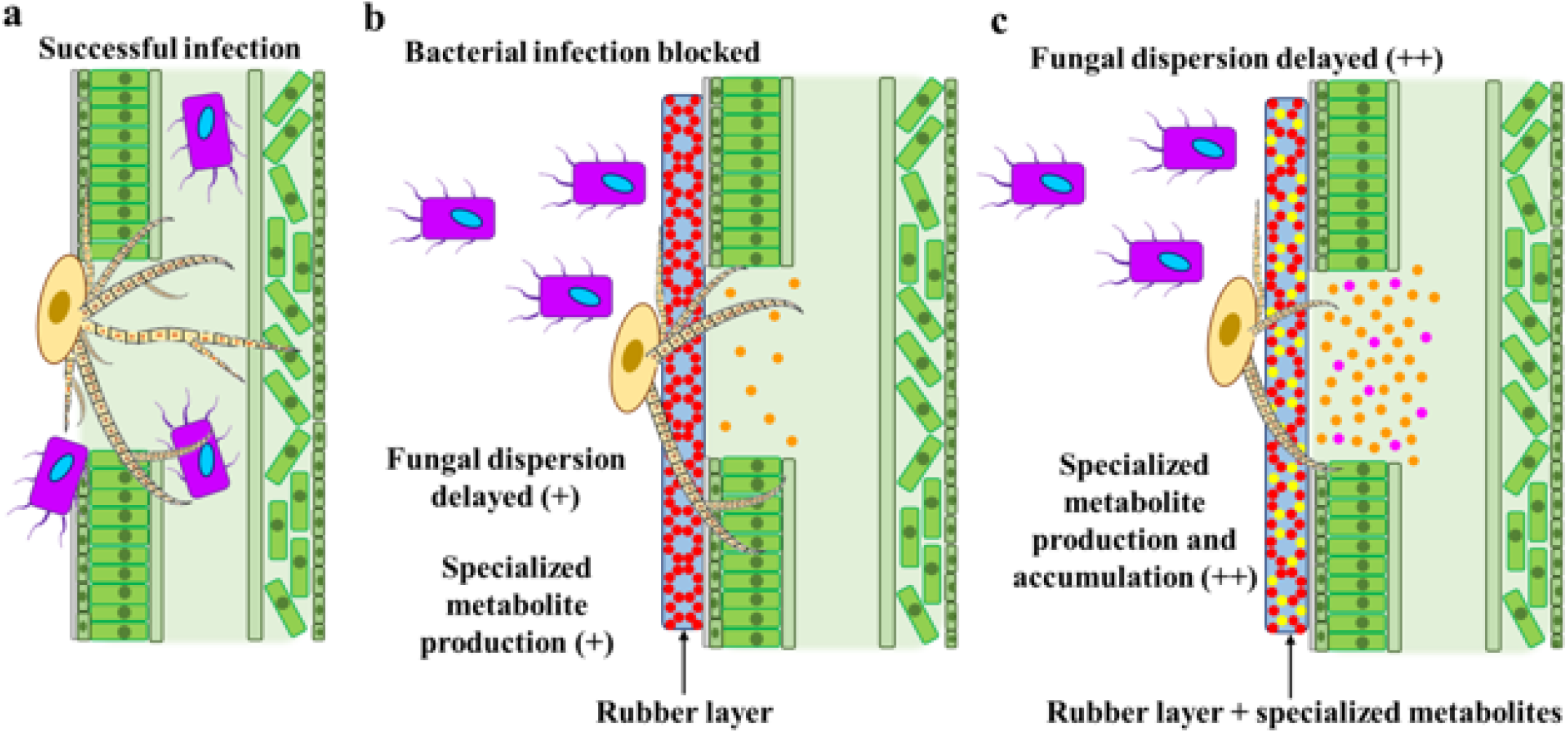
Chemo-mechanical defense system in latex against general fungal and bacterial pathogens. A, Open wound in leaf tissue. Pathogenic microorganisms are free to enter and cause potential infections. B, Plant wound covered with a rubber latex after latex coagulation. The mechanical barrier is sufficient to avoid bacterial infections, but only slows fungal spread. C, Plant wound covered with a polymerized layer of rubber after latex coagulation is reinforced with specialized metabolites. The fungal pathogens grow and disperse at a lower rate, while the induced chemical defense system is activated in the laticifer.

## Conclusion

The metabolome of latex is differentiated from other tissues in the same plant, which denotes their chemical specialization. Moreover, the variation of the chemical profiles of latexes is much lower when compared with those of other plant tissues. This stability is correlated with polyisoprenes and triterpenoids. This evidences latex as a constitutive defense system that is not substantially influenced by environmental factors. In this system, the major constituents function as an instantaneous multi-pronged defense system in plants. The polyisoprenes firstly act as a mechanical barrier to stop and/or retard infections and/or herbivory, while triterpenoids fight residual microorganisms. Therefore, the metabolic constraint of constitutive plant latexes is a key strategy for their success on fighting against herbivores and pathogens.

## Materials and methods

### Plant material and collection

Leaves, roots, and latexes of *Euphorbia glareosa* Pall. Ex M. Bieb., *Euphorbia amygdaloides* L., and *Euphorbia palustris* L. were collected in several locations in Serbia: Deliblatska peščara, Zagajička brda, Titelski breg, Avala, Mali Jastrebac, Kosmaj, Borča, Čenta, and Šajkaš at 44°56’41.44″N 21°4’29.52″E, 44°55’48.29″N 21°11’51.68″E, 45°13’23.92″N 20°13’45.03"E, 44°41’11.41″N 20°30’53.20"E, 43°23’3.51″N 21°36’47.97"E, 44°28’28.45″N 20°34’28.10"E, 44°54’48.34″N 20°26’32.51"E, 45°5’58.15″N 20°22’42.06"E, and 45°15’13.14″N 20°6’31.21"E, respectively, in June, 2017. The plant materials were identified by Pedja Janaćković, and voucher specimens were deposited at the Herbarium of the Botanical Garden “Jevremovac” University of Belgrade, Belgrade, Serbia (voucher numbers: *Euphorbia glareosa* (BEOU17303), *Euphorbia amygdaloides* (BEOU17306), and *Euphorbia palustris* (BEOU17304)). Latex samples were obtained by making incisions in the plant stems with a sterile razor blade and collecting approximately 1 mL of latex in a sterile 2 mL-microtube containing 400 μL of MeOH. The samples were stored at −20 °C until they were freeze-dried. Leaf and root samples were manually collected from the plant and placed into plastic hermetic bags with silica gel. This material was stored at −20 °C until processed. The leaf and root samples were processed by grinding with liquid nitrogen and then freeze-drying. Dry leaf (10 g) and root (5 g) powders were extracted with methanol with sonication during 15 min. The solvent was evaporated with a rotary evaporator, and the extracts were taken to total dryness with a speed-vacuum dryer.

### Insects, fungi and bacteria

Larvae of *Mamestra brassicae* were kindly provided by Pieter Rouweler from the Entomology Department at Wageningen University. *Pseudomonas putida* (NCCB26044) was purchased from The Netherlands Culture Collection of Bacteria. *Pseudomonas fluorescens* and *Pseudomonas viridiflava* were kindly provided by Dr. Paolina Garbeva (Kurm et al., 2019). *Alternaria alternata* (CBS 102.47) strain was purchased from the collection of the Westerdijk Fungal Biodiversity Institute, and *Botrytis cinerea* was kindly provided by Dr. Jan van Kan (Van Kan et al., 2017).

### HPTLC-DART-MS analysis

For thin-layer chromatography, latex extracts were diluted with methanol to a final concentration of 2 mg/mL. An automatic TLC sampler (ATS 4) (CAMAG, Muttenz, Switzerland) with a 25 μL Hamilton syringe was used to apply 30 μg of all of the samples as 6 mm bands on 20 × 10 cm HPTLC silica gel plates (60 F254) (Merck). The samples were applied at 20 mm from the lateral edges and 10 mm from the bottom of the plate. The distance between bands was 10 mm, resulting in 18 tracks per plate. Chromatographic development was performed in an automatic developer (ADC2) (CAMAG, Muttenz, Switzerland). The samples were separated with a mixture of toluene–ethyl acetate (8:2, v/v). The saturation time was 20 min, and humidity was set to 37% using a saturated MgCl_2_ solution. The solvent migration distance was 75 mm from the application point. The HPTLC system was controlled by Vision Cats software.

For HPTLC-DART-MS analysis, each track of the plate was cut in 5 mm-wide strips using a smart glass plate cutter (CAMAG, Muttenz, Switzerland). The HPTLC strips were individually placed on a motorized rail to be moved to the ionization region. The plates were ionized with a DART ion source (Ion Sense, Tokyo, Japan) using helium gas (purity of 99.999%) at 450 °C and 3 L h^−1^. The plate scan speed was 0.2 mm s^−1^, and it was controlled with DART control software (Ion-Sense). The distance from the ion source to the plate was 1.5 cm. Detection was performed with an AccuTOF-TLC (JEOL, Tokyo, Japan) in positive ion mode. The TOF-MS was set with a peak voltage of 800 V and a detector voltage of 1,900 V.

### ^1^H NMR analysis

Five mg of freeze-dried latexes were re-suspended in 1 mL of CH_3_OH-*d_4_* containing 3.93 mM hexamethyldisiloxane (HMDSO) as the internal standard, and ultrasonicated for 20 min. For leaf and root samples, 5 mg of the MeOH extract were dissolved in 1 mL of CH_3_OH-*d_4_*. All of the solutions were centrifuged at 13000 rpm for 10 min, and 300 μL of the supernatant were transferred into 3 mm-NMR tubes. The ^1^H NMR analysis was carried out with an AV-600 MHz NMR spectrometer (Bruker, Karlsruhe, Germany), operating at a proton NMR frequency of 600.13 MHz. For internal locking, CH_3_OH-*d_4_* was used. All ^1^H NMR consisted of 64 scans requiring 10 min and 26 s as acquisition time using the parameters of 0.16 Hz/point, pulse width (PW) = 30° (11.3 μs), and relaxation time of 1.5 s. A pre-saturation sequence was used to suppress the residual water signal using low power selective irradiation at H_2_O frequency during recycle delay. The FIDs were Fourier transformed with exponential line broadening of 0.3 Hz. The resultant spectrums were manually phased and baseline corrected, and calibrated to HMDSO at 0.06 ppm using TOPSPIN V. 3.0 (Bruker).

### Gas chromatography coupled to mass spectrometry (GC-MS) analysis

Dried latexes (5 mg) were extracted with 1 mL of chloroform. The extract was taken to total dryness with a speed-vacuum dryer. The dried extracts were re-dissolved with 100 μL of pyridine by ultrasonication for 5 min. To this, 100 μL of BSTFA + TMCS (99:1, Supelco) were added, and the solutions were heated at 80 °C for 50 min. The solutions were then centrifuged at 13000 rpm for 10 min, and the supernatants were transferred to micro-inserts for GCMS analysis on a 7890A gas chromatograph equipped with a 7693 automatic sampler coupled to a 5975C mass single-quadrupole detector (Agilent, Folsom, CA, U.S.A.). Separation was performed on a DB5 GC column (30 m x 0.25 mm, 0.25 μm thickness, JW Science, Folsom, CA, U.S.A.) with helium (99.9% purity) as the carrier gas at a flow rate of 1 mL/min. The initial oven temperature was 100 °C for 2 min, and then ramped at 10 °C/min to 270 °C, held for 1 min, ramped again to 290 °C at 5 °C/min for 15 min, and then to 300 °C at 5 °C/min and held for 3 min. The injector was set at 280 °C, and 1 μL of each sample was injected in splitless mode. The interface temperature was 280 °C, and the ion source and quadrupole temperature of the mass detector was 230 °C and 150 °C, respectively. Ionization energy in EI mode was 70 eV, and peaks were identified by comparison of the ion spectra with the NIST library (version 2008).

### Liquid chromatography coupled to mass spectrometry (LC-MS) analysis

Five milligrams of each latex sample were individually re-suspended in 1 mL of a methanol-water solution (80:20, v/v) and ultrasonicated for 15 min. The resulting extracts were diluted in a 1:10 (v/v) ratio to a final concentration of 0.5 mg mL^−1^. The samples were filtered with 0.20 μm membrane filters and analyzed using an Acquity UPLC HSS T3 column (2.1 mm × 100 mm, 1.7 μm; Waters). Samples were eluted with a gradient of 0.1% formic acid in water (A) and 0.1% formic acid in acetonitrile (B) starting at 10% B (0 – 30 min), 100% B (30 – 35 min), and 10% B (35 – 40 min) at 40 °C at a flow rate of 0.3 mL min^−1^. MS detection was performed on a QTOF mass spectrometer (Bruker Impact HD) equipped with an electro-spray ionization (ESI) source. The capillary voltage was 4000 V, and the drying temperature was 350 °C at 6 L min^−1^. The samples were analyzed in negative and positive mode in the range of 50-1200 *m/z*. Data acquisition, alignment, peak picking, and neutral losses calculations were performed using Progenies QI software version 2.3 (Nonlinear Dynamics: a Waters Company, Newcastle, UK). Quality control (QC) samples consisted of a blend of all samples that was injected every five samples. Extraction solvent was injected as a blank. The data were normalized to total intensity, and filtered by deleting mass features detected in blank samples at higher response levels than in latex samples. Data filtering resulted in 8015 mass features for data acquired in positive mode and 2064 features in negative mode. Compounds were identified by comparison of their exact mass with the Dictionary of Natural Products. A threshold of 10 ppm was set as the mass error for possible matches.

### *cis*-1,4-Polyisoprene (rubber) solution preparation

Five grams of rubber (Sigma-Aldrich) were cut into small pieces and placed in 100 mL of chloroform. The rubber pieces were left in the solvent for 3 h to swell, and then manually stirred every 20 min for 1 min until a semi-clear solution was obtained. The volume was then adjusted to 140 mL, and stirred until a clear rubber slurry was formed.

### Microdilution antibacterial assay

The broth microdilution method was utilized to determine the minimal inhibitory concentration (MIC) of the tested triterpenes according to the Clinical Laboratory Standards Institute guideline. The strains were inoculated on Mueller-Hinton agar (MHA) plates and incubated overnight at 37 °C. From the overnight cultures, a single colony was used to inoculate 10 mL of Mueller-Hinton broth (MHB) and incubated at 37 °C under constant agitation (150 rpm). The bacterial suspensions were further adjusted with the addition of MHB to 0.5 of turbidity of the McFarland scale (10^6^ CFU/mL). In parallel, the compounds were dissolved in DMSO and diluted to reach final concentrations in the well from 512 μg/mL to 16 μg/mL, and then taken to a volume of 100 μl in each well with MHB. Each well was then inoculated with 50 μl of the 0.5 McFarland bacterial suspensions, and incubated for 24 h at 30 °C. The final concentration of DMSO in the well was 5%, which was also used as a negative control. A 100 μg/mL solution of spectinomycin in one well was used as a positive control. The bacterial growth was measured by optical density at 600 nm in a well microtiter plate reader (SPARK 10M, TECAN). The MIC value was defined as the lowest concentration of a compound that completely inhibited the bacterial growth at 24 h. All experiments were performed in triplicate. For *P. viridiflava*, the inoculations were done in nutrient agar 2 plates, and the assays were carried out in nutrient broth 2 at 28 °C.

### Antibacterial assay

The bacterial strains were prepared in the same manner as in the microdilution method. The agar plates with treatments were prepared by filling 45 mm Petri dishes with 7 mL of MHA (nutrient agar 2 for *P. viridiflava*). After medium solidification, 3 mL of rubber solution was poured over the medium and left to dry for 2 h in a fume hood, resulting in a homogeneous rubber layer of ca. 0.05 mm of thickness. Finally, the plates were exposed for 10 min to UV light for sanitation. The negative controls were MHA plates without rubber covering. Four replicates were performed for treatments and controls of *Pseudomonas putida*, *Pseudomonas fluorescens*, and *Pseudomonas viridiflava*. The bacterial growth was evaluated at 24 h and 48 h. To determine whether bacteria was unable to pass through the rubber layer to the agar, the rubber layer from the inoculated treatment plates was removed, and the plates were incubated for 24 h at 28 ± 2 °C. Plates were then inspected for the appearance of bacterial colonies at the inoculation points.

### Antifungal assays

The treated plates were prepared similarly to those used for antibacterial assays, but using potato dextrose agar (PDA). Seven-millimeter agar plugs with *Alternaria alternata* and *Botrytis cinerea* were placed on top of the rubber layer of the treated plates. The negative controls were PDA plates without a rubber layer, PDA plates just with rubber solution, and PDA plates with a combination of rubber solution and latex powder. To observe if the fungi grew over the rubber layer or penetrated it, the layer was manually removed from the plate. Four replicates were performed for each fungal strain, and their growth and colony diameter were measured at 48 h. The incubation temperature was 26 °C. In the case of the rubber and latex combination, *E. palustris* and *E. myrsinites* were chosen as models due to their very different triterpenoid profile. Five milligrams of all samples of each species were mixed to obtain a representative sample of each latex. The latex was then dissolved in the rubber solution to reach a concentration of 200 μg/mL. The plates were dried in the same way as in previous experiments.

### Minimum inhibitory concentration (MIC)

24-methylenecycloartenol and β*-*sitosterol, as representatives of different steps of the phytosterol pathway, and α*-*amyrin, β*-*amyrin, ursolic and oleanolic acids, as representatives of different steps in the triterpene pathway, were selected to be tested against *Botrytis cinerea*. The compounds were dissolved in methanol, and tested in two-fold dilution series from 2000 μg/mL until 62.5 μg/mL. The spore solution was adjusted to 2.5 x 10^5^ spores/mL in the well, and the final concentration of methanol in the well was 5%. The plates were sealed with a parafilm layer and incubated at 26 °C. The positive control consisted of nystatin in the same range of concentrations, and the negative control consisted of media with a final concentration of 5% of methanol in the well. The MIC value was defined as the minimum concentration in which there was visible total inhibition of fungal growth at 16 h. The plates were examined under a stereoscopic microscope at 16 h intervals to observe any further effects on fungal growth.

### Anti-herbivore activity

The diet was prepared by mixing 28 g of agar, 160 g of cornflower, 50 g of beer-yeast, 2 g of sorbic acid, 1.6 g of methyl-4-hydroxybenzoate, 8 g of ascorbic acid, and 0.1 g of streptomycin per litre of water. The ingredients were added to warm water with continual stirring. The diet (15 mL) was placed in individual plastic containers and left to solidify at room temperature. For treatments, the dry methanol extracts were re-suspended in 2 mL of ultrapure water and ultrasonicated for 10 min (2X). These were added to the diets while they were still semiliquid, manually mixed, and left to solidify at room temperature. The treatments consisted of methanol extracts of latexes, leaves, and roots. From the 10 samples from each location of each species, three random samples were mixed to form one composed sample. Therefore, three composed samples from each region of each plant species were obtained for the three tissues, resulting in nine replicates for each species of each tissue and three replicates for different locations. The final concentration of all of the treatments was 200 μg/mL. The negative control consisted of ultrapure water, and the positive control was a 200 μg/mL concentration of abamectin in the diet. The weight of the larvae was recorded after 5 d, and the weight of treated and untreated larvae (negative control) were compared.

### Data processing and multivariate data analysis

The NMR spectra were bucketed using AMIX 3.9.12 (Bruker BioSpin GmbH, Rheinstetten, Germany). Bucket data were obtained by spectra integration at 0.04 ppm intervals. Peak intensity of individual peaks was scaled to total intensity, and recorded from *δ* 0.20 to 10.02. Because of the residual signals of D_2_O and CH_3_OH-*d_4_*, regions of δ 4.75 – 4.9 and δ 3.28 – 3.34 were excluded from the analysis, respectively. Multivariate data analysis was performed using SIMCA P (v.15, Umeå, Sweden). Principal component analysis (PCA) and partial least square discriminant analysis (PLS-DA) were conducted for ^1^H NMR and LC-MS data. For PCA and PLS-DA analyses, the data was scaled using the unit variance (UV) method.

To assess the effect of the geographical region on chemical diversity, a soft independent model of class analogy (SIMCA) analysis was performed using geographical origin and plant species as PCA-classes separately in each plant species. The distances to the model (DModx) values were calculated setting each plant species as a PCA-class in the species effects model, and the same geographical region in each species for the geographical origin effect in tissues. Data were scaled using the UV method, and the DModx values were transformed to their corresponding logarithm values. The logarithmic averaged DModx values (n = 30) ± standard error of each model were used as a measure of the strength of each factor in the chemical homogeneity of the samples.

In order to obtain and compare the total correlation of the effects of the geographical origin of the species on the chemical composition of the samples, PLS-DA modelling using UV scaling was also performed on each individual data set. The averaged *Q^2^* from the permutation test (100 permutations) was used as a measure of the strength of the effects of the species and geographical origin on the chemical profile differences among the different plant tissues.

For the antifungal bioassays, the radial growth of the treatments was compared to their corresponding control by a Dunnett test setting the control sample as a control for the comparison of the treatments to the control, and setting the rubber treatment as a control for its comparison to the latex supplemented treatments (*α =* 0.05) for *Alternaria alternata*, and after log transformation of the data for *Botrytis cinerea*. The anti-herbivory activity data variance, homogeneity, and mean comparison were done with a type 2 ANOVA, and the mean comparison was performed with a least-square means test (*α =* 0.05) after log transformation using R software (V 1.1.456).

## Supplemental Data

The following supplemental materials are available.

Supplemental Figure S1. Principal component analysis of latex, leaves, and roots of three *Euphorbia* species collected at different locations of Serbia. A, latex, B, leaves, C, roots. 1, *Euphorbia amygdaloides*, 2, *Euphorbia glareosa*, 3, *Euphorbia palustris*. The separation of the samples by geographical origin is much more evident in leaves and much less evident in latexes.

**S**upplemental Figure S2. Effect of triterpenes over the growth of *Botrytis cinerea.* A, Nitastine [2 mg/mL], B, Nitastine [62.5 μg/mL], C, 24-methylenecycloartanol [2 mg/mL], D, 24-methylenecycloartanol [62.5 μg/mL], E, *α*-amyrin [2 mg/mL], F, *α*-amyrin [62.5 μg/mL], G, *β*-amyrin [2 mg/mL], H, *β*-amyrin [62.5 μg/mL], I, oleanoic acid [2 mg/mL], J, oleanoic [62.5 μg/mL], K, ursolic acid [2 mg/mL], L, ursolic acid [62.5 μg/mL], M, *β*-sitosterol [2 mg/mL], N, *β*-sitosterol [62.5 μg/mL], O, negative control.

## ACKNOWLEDGEMENTS

The first author expresses gratitude to the Mexican Scientific Council (CONACyT) for supporting his Ph.D. scholarship (No. 410812).

